# Pyridox(am)ine 5’-phosphate oxidase deficiency induces seizures in *Drosophila melanogaster*

**DOI:** 10.1101/580050

**Authors:** Wanhao Chi, Atulya Iyengar, Monique Albersen, Marjolein Bosma, Nanda M. Verhoeven-Duif, Chun-Fang Wu, Xiaoxi Zhuang

## Abstract

Pyridox(am)ine 5’-phosphate oxidase (PNPO) is a rate-limiting enzyme in converting dietary vitamin B6 (VB6) to pyridoxal 5’-phosphate (PLP), the biologically active form of VB6, and involved in the synthesis of neurotransmitters including GABA, dopamine, and serotonin. In humans, PNPO mutations have been increasingly identified in neonatal epileptic encephalopathy and more recently also in early-onset epilepsy. Till now, little is known about the neurobiological mechanisms underlying PNPO-deficiency-induced seizures due to the lack of animal models. Previously we identified a c.95 C > A missense mutation in *sgll* - the *Drosophila* homolog of human *PNPO* (*hPNPO*) and found mutant (*sgll*^95^) flies exhibiting a lethal phenotype on a diet devoid of VB6. Here we report the establishment of both *sgll*^95^ and ubiquitous *sgll* knockdown (KD) flies as valid animal models of PNPO-deficiency-induced epilepsy. Both *sgll*^95^ and *sgll* KD flies exhibit spontaneous seizures before they die. Electrophysiological recordings reveal that seizures caused by PNPO deficiency have characteristics similar to that in flies treated with GABA antagonist picrotoxin. Both seizures and lethality are associated with low PLP levels and can be rescued by ubiquitous expression of wild-type *sgll* or *hPNPO*, suggesting the functional conservation of the PNPO enzyme between humans and flies. Results from cell type-specific *sgll* KD further demonstrate that PNPO in the brain is necessary for seizure prevention and survival. Our establishment of the first animal model of PNPO deficiency will lead to better understanding of VB6 biology, the *PNPO* gene and its mutations discovered in patients, and can be a cost-effective system to test therapeutic strategies.

## Introduction

Mutations in an autosomal gene encoding pyridox(am)ine 5’-phosphate oxidase (PNPO) cause neonatal epileptic encephalopathy (NEE, OMIM #610090), a severe neurological disease that leads to death if untreated [1]. Furthermore, PNPO mutations have more recently been identified also in patients with infantile spasms [2] and early-onset epilepsy [3, 4, 5]. While the causal relationship between PNPO deficiency and NEE has been established on the molecular level since 2005 [1], the neurobiological mechanisms of how PNPO deficiency leads to NEE and related epilepsy syndromes remain to be established.

PNPO is a rate-limiting enzyme in the synthesis of vitamin B6 (VB6), which comprises a group of six different forms including pyridoxine (PN), pyridoxal (PL), pyridoxamine (PM), pyridoxine 5’-phosphate (PNP), pyridoxal 5’-phosphate (PLP), and pyridoxamine 5’-phosphate (PMP) [6, 7]. PNPO converts PNP and PMP to PLP, the biologically active form of VB6, which is a co-factor for more than 140 different enzymes that are involved not only in the metabolism of amino acids, glycogen, and unsaturated fatty acids but also in the synthesis of various neurotransmitters including *γ*-aminobutyric acid (GABA), dopamine, and serotonin [8]. PNPO activity is essential to humans and other animals because unlike plants, fungi, and bacteria, they cannot synthesize VB6 *de novo* [9]. PNPO is highly expressed in liver, brain, and kidney in mammals [10] and previous studies have implicated the liver [11] as the primary site of conversion of dietary VB6 to PLP, which is further transported by circulation to different tissues/organs, including the brain [12, 13]. However, the contribution of brain-expressed PNPO to seizures and lethality, is unknown.

In recent years, PNPO mutations have been increasingly reported in human patients [1, 3, 5, 14, 15, 16, 17, 18, 19, 20, 21, 22, 23, 24, 25, 26, 27, 28]. Seizures in PNPO deficient patients usually do not respond to conventional anti-epileptic drugs but is responsive exclusively to PLP, or to PN in patients with partial PNPO deficiency [17, 20, 23]. Despite its effectiveness in seizure control, PLP or PN treatment has little effect on other neurological defects observed in PNPO deficiency patients, such as developmental delay or intellectual disability [20, 24, 26]. Moreover, long-term PLP treatment causes hepatic cirrhosis [29, 30]. There is a need for valid animal models to study the fundamental biology of PNPO and VB6, their role in neurodevelopment and neurotransmission, and to explore other treatment options for PNPO deficiency.

Based on the nutritional conditional lethal phenotype, we have recently identified a *Drosophila melanogaster* homolog gene of human *PNPO* (*sgll*) and identified a mutant with partial PNPO deficiency (*sgll*^95^) [31]. The *sgll*^95^ flies survive well on a normal diet, but due to their low PNPO activity, they exhibit lethality when reared on a sugar-only diet (4%sucrose in 1% agar, i.e., diet devoid of VB6). This nutritional conditional lethal phenotype can be rescued by the supplementation of either PN or PLP. Here we report that human and *Drosophila* PNPOs are functionally conserved and that PNPO deficiency in *Drosophila* causes seizures that are defined by both behavioral and electrophysiological parameters. These PNPO-deficiency-induced-phenotypes are associated with low levels of PLP and PNPO deficiency restricted to the neural tissue is sufficient to cause seizures and lethality.

## Results

### *sgll*^95^ flies exhibit seizure-like behavior before they die

Consistent with our previous report [31], we found that *sgll*^95^ homozygotes reared on a sugar-only diet, consisting of 4% sucrose in 1% agar, exhibited a striking lethality phenotype compared to *w*^1118^ counterparts. As shown in Figure 1A, nearly all *sgll*^95^ mutants had died after four days while no control flies had died. Before death (2 - 3 days on the sugar-only diet, green box in Figure 1A), *sgll*^95^ mutants displayed bouts of rapid ‘wing-buzzing’, and ‘body-rolling’, and later on, they often exhibited a ‘wings-up’ posture (Figure 1B). These seizure-like behavioral phenotypes occurred spontaneously and did not appear to be initiated by mechanical shock [32, 33, 34, 35] or temperature stress [36, 37, 38, 39, 40, 41].

**Figure 1.**
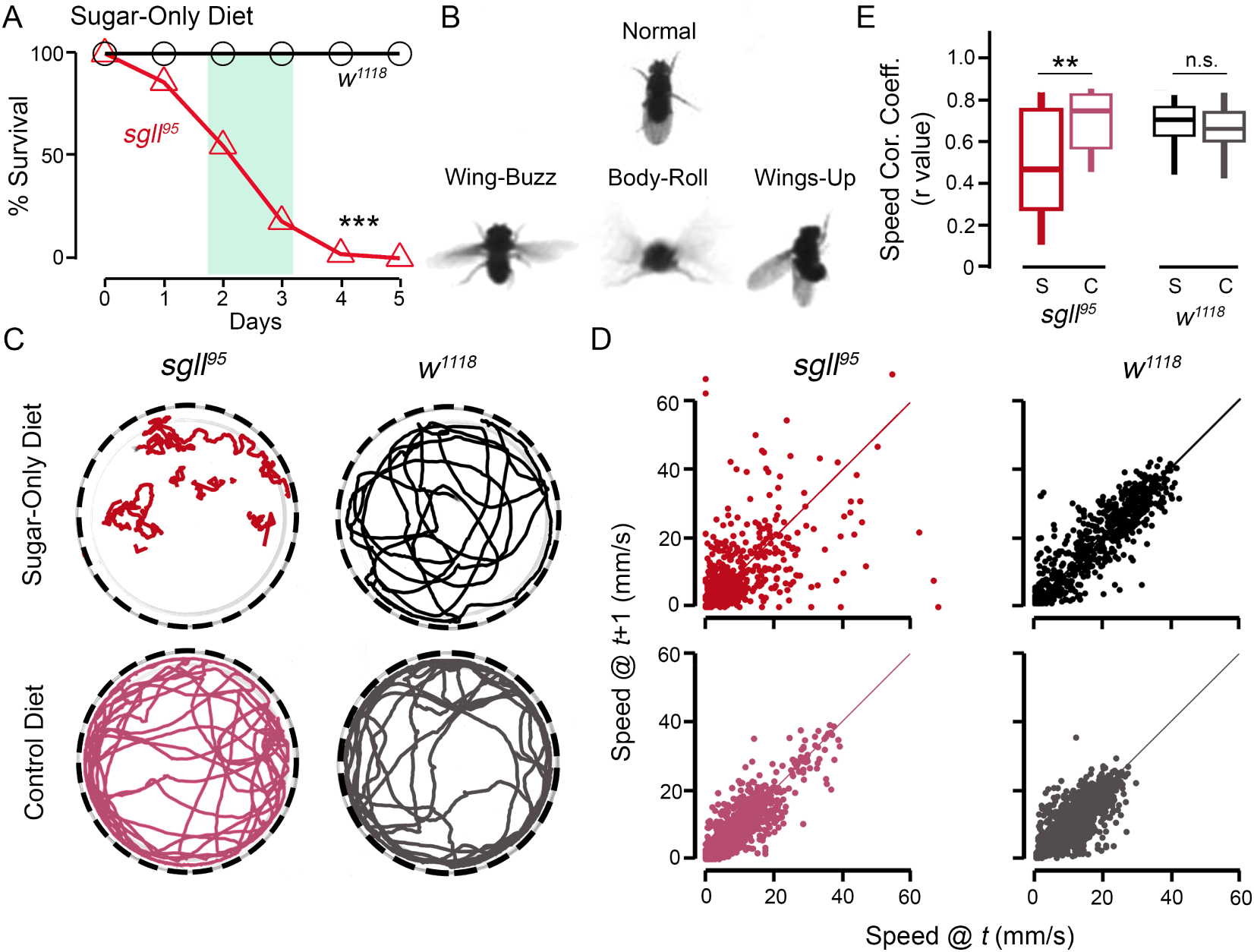
*sgll*^95^ flies show seizure-like behavior before they die when reared on the sugar-only diet. A) Survival of *sgll*^95^ and *w*^1118^ on the sugar-only diet. *** *P* < 0.001, Log-rank test, n = 100-101. The green block indicates the time window when flies were characterized for the seizure phenotypes. B) Representative images of normal and seizure-like behaviors from video recordings. C) Representative travel traces from video recordings. Each trace is from one fly. The fragmented trace from the *sgll*^95^ fly on the sugar-only diet is due to the faster movement of the fly than the frame window (50 ms). D) Speed correlation plots between the frame *t* and frame *t*+1. Each plot is corresponding to the one in panel C. E) Summary plot of the speed correlation coefficient (Speed Cor. Coeff.). S: sugar-only diet, C: normal diet, n.s.: *P* > 0.05, ** *P* < 0.01, Two-way ANOVA with Tukey’s *post hoc*, n = 14-22 per genotype per condition.

To examine these behaviors, we recorded individual *sgll*^95^ or *w*^1118^ control flies walking in a 60 mm-wide open arena with a high-resolution video camera for 3 min (See Methods for details). Fly positions were then tracked with the IowaFLI Tracker [42], and the corresponding travel traces from each fly were plotted. We observed a smooth travel trace from each *w*^1118^ control fly (n = 22 on sugar; n = 19 on the normal diet) or from *sgll*^95^ fly on the normal diet (n = 14). However, travel traces of *sgll*^95^ flies reared on the sugar-only diet (n = 21) were heterogeneous with some traces similar to that of controls whereas others did not show smooth trajectories at all. Representative traces are shown in Figure 1C.

To quantify these travel traces, we plotted each trace on a scatter plot where the speeds of a fly during a frame *t* are plotted against the speeds in the next frame *t*+1 (Figure 1D) and then we calculated the speed correlation coefficients. The reasoning is that a smooth travel trace indicates consistent locomotion, i.e., similar speeds at two consecutive time points. On the scatter plot, all points from a smooth trace would land on the lower left to upper right diagonal line and give a high correlation coefficient of speed. On the opposite side, the less smooth travel traces will have their points spread out from the diagonal line and give lower correlation coefficients. Indeed, the average correlation coefficient is 0.654 ± 0.116 (Mean ± SD), 0.672 ± 0.120, 0.701 ±0.140, and 0.493 ± 0.260 for *w*^1118^ on the normal diet, *w*^1118^ on the sugar-only diet, *sgll*^95^ on the normal diet, and *sgll*^95^ on the sugar-only diet, respectively. Therefore, consistent with qualitative distinctions between *sgll*^95^ and *w*^1118^ flies, we found significantly reduced speed correlation coefficients in *sgll*^95^ when reared on the sugar-only diet (Figure 1E, *P* = 0.0047, gene by condition interaction *P* = 0.0065).

### Ubiquitous RNAi-mediated knockdown of *sgll* leads to lethality and seizure-like behavior when reared on the sugar-only diet

We have previously shown that ubiquitous *sgll* knockdown (KD, i.e. *actin*-Gal4/UAS-*sgll* RNAi) flies also show the conditional lethal phenotype, which can be rescued by PLP but not PN [31], indicating severe PNPO deficiency in these flies compared to that in *sgll*^95^ flies. Indeed, *sgll* KD flies survived shorter than *sgll*^95^ flies (compared males to males); all *sgll* KD flies died within four days (Supplemental Figure S1A) while all *sgll*^95^ flies died within five days (Figure 1A). Prior to death, *sgll* KD flies displayed seizure-like behavior similar to that in *sgll*^95^ flies including the wing buzzes, wings-up posture and convoluted walking trajectories (Figure S1 B&C). The average speed correlation coefficient of *sgll* KD flies is 0.423 ± 0.240 (Mean ± SD), which is significantly lower than that from the parental controls (0.644 ± 0.139, and 0.645 ± 0.233 for *actin*-Gal4/+ and UAS-*sgll* RNAi/+, respectively. Figure S1 D; *P* = 5.947e-05).

### Spontaneous seizure discharges in *sgll*^95^ and *sgll* KD mutants reared on the sugar-only diet

To monitor electrophysiological activity and identify aberrant patterns in *sgll* mutants, we undertook an analysis of flight muscle activity in a tethered fly preparation (Supplemental Video 1) [43, 44]. The isometric contractions of the dorsal longitudinal muscle (DLM) enable prolonged recordings of spiking activity with minimal damage to the muscle. Several studies have taken advantage of this stable recording preparation to monitor aberrant spike discharges associated with seizures that occur spontaneously in mutant flies [40, 41], seizures that are triggered by high-frequency electroconvulsive stimulation [43, 45, 46] or by proconvulsant administration [47]. Importantly, the bouts of DLM spiking activity during seizure discharges appears to be temporally correlated with extracellular synchronous activity across different body-axes [46].

We recorded spontaneous DLM spiking activity not associated with flight from *sgll*^95^, *sgll* KD flies and *w*^1118^ control flies, reared on either the normal diet or the sugar-only diet (Figure 2A). Flies reared on the normal diet displayed short bouts of a few spikes associated with wing depression events during grooming activity (Supplemental Video 1, also see reference [47]). Over the recording session (∼240 s) the average firing frequency (total # spikes / recording duration) was ∼0.3 Hz, for the three genotypes tested (Figure 2B). In a striking contrast, recordings from both *sgll*^95^ and *sgll* KD flies reared on the sugar-only diet showed high-frequency spike burst discharges qualitatively distinct from the grooming-related spiking seen in their counterparts reared on the normal diet (Figure 2A), and there was also significantly more firing (median firing rate of: 2.03 Hz.- vs. 0.32 Hz.- for *sgll*^95^, and 7.91 vs. 0.07 Hz for *sgll* KD flies, *P* = 0.0045 and *P* = 1.138e-6, for *sgll*^95^ and *sgll* KD, respectively. Figure 2B). These spike bursts occurred simultaneously in the left and right muscles (data not shown), suggesting that they were triggered by relatively widespread events across the nervous system. Notably, our sample included *sgll*^95^ and *sgll* KD flies with ‘wings-up’ as well as normal wing posture (Figure 1B). We found that between our recordings of *sgll*^95^ and *sgll* KD flies, all wings-up flies displayed spike bursts (6 / 6), and several individuals with normal wing posture did that as well (5 / 8 flies), suggesting that the burst discharge phenotype likely initiates prior to appearance of the ‘wings-up’ posture.

**Figure 2.**
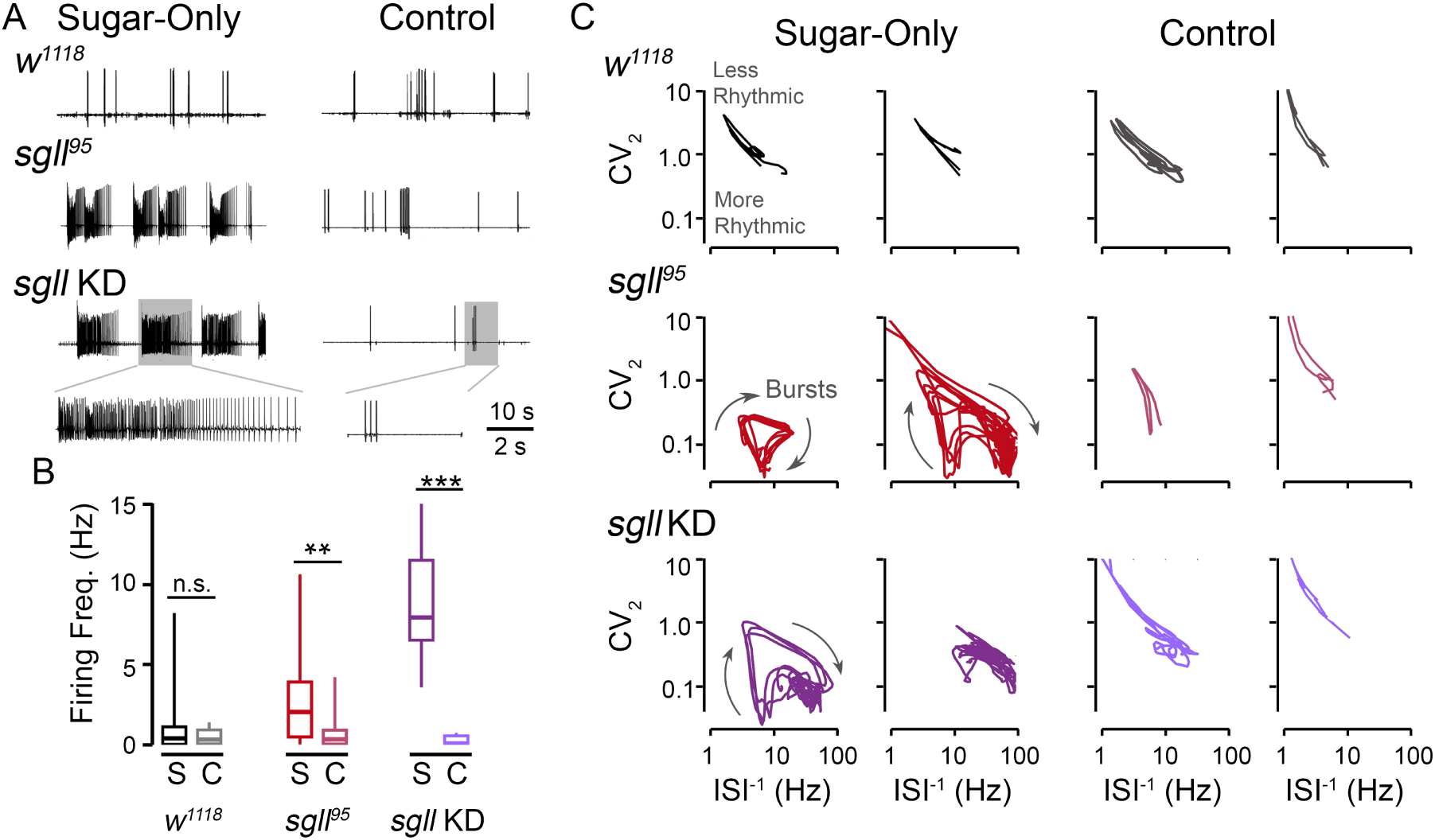
Spontaneous spike discharges in flight muscles of *sgll* mutants. A) Representative traces of DLM spiking in *w*^1118^ control flies, *sgll*^95^ mutants, and ubiquitous RNAi knockdown of *sgll* by *actin*-Gal4 driver (i.e. UAS-*sgll* RNAi/*actin*-Gal4, indicated as *sgll* KD). Flies were reared on the sugar-only diet (Sugar-only, left column) or normal diet (Control, right column). Note the difference between the high-frequency spike discharges in the *sgll*^95^ and *sgll* KD traces and the spiking in *w*^1118^ flies and *sgll* mutants on the normal diet (see expanded traces). The spiking in flies on the normal diet is associated with grooming behavior (Supplemental Video 1, also see reference (43)). B) Summary plot of the average firing frequency over the duration of the recordings (∼90 s) in the respective genotypes on 4% sucrose (S) or normal (C) diet. n.s.: *P* > 0.05, ** *P* < 0.01, *** *P* < 0.001, Rank-sum test. n = 7-11 flies. (C) Plots of the trajectory of the instantaneous firing frequency (ISI^−1^) vs. the instantaneous coefficient of variation (CV2) during DLM spiking. For each genotype/condition, two representative trajectories are shown. The oscillations indicated in *sgll*^95^and *sgll* KD trajectories correspond to individual burst discharges in the DLM trace.

An important characteristic of the DLM spike trains recorded was the highly variable nature of inter-spike intervals (ISIs). Within a spike train, the instantaneous firing frequency between two spikes (defined as the inverse of the ISI, i.e. ISI^−1^) could range from less than 1 Hz to more than 100 Hz within a bout. To quantitatively delineate the grooming-related spike patterns, we employed a phase-space analysis of ISIs [47]. For the sequence of spikes within the recording, we plotted the ISI^−1^ of each spike against the spike’s instantaneous coefficient of variation, CV_2_, a measure of variability between successive ISI^−1^ values (see Methods for computational details). The resulting ISI^−1^ vs CV_2_ trajectories have been shown to clearly distinguish between spike patterns associated with flight, grooming, and electroconvulsive stimulation-induced seizure discharges in *Drosophila* [47].

The ISI^−1^ vs. CV_2_ trajectories of the burst discharges in *sgll* mutants on the sugar-only diet were readily distinguished from the spiking activities in counterparts on the normal diet and in *w*^1118^ flies (representative trajectories shown in Figure 2C). Specifically, trajectories of *sgll*^95^ flies on the sugar-only diet display ‘loops’ consisting of the ISI^−1^ of the trajectory with abrupt acceleration (often peaking between 50 and 100 Hz) followed by a gradual deceleration characterized by relatively low CV_2_ values. A few *sgll* KD flies displayed more extreme trajectories which were more compact, reflecting continuous high-frequency firing. In contrast, the ISI^−1^ vs. CV_2_ trajectories of *sgll* mutants on the normal diet as well as that of *w*^1118^ flies generally ambled within a limited region of higher CV_2_ values (> 0.5) reflecting a high degree of variability in successive spike intervals, consistent with observations of grooming spike patterns in other wild-type fly strains [47].

### Seizures and lethality are correlated with low levels of internal PLP

PNPO converts intracellular PNP and PMP to PLP (Figure 3A), and PNPO deficiency is expected to result in a low level of PLP. However, both normal and decreased PLP levels have been reported in human PNPO deficient patients [26, 48]. Such inconsistency is conceivably due to confounding factors such as heterogeneous genetic backgrounds and different diets [3]. Since both genetic background and dietary conditions were well controlled in flies, we reasoned that by comparing the PLP level between *sgll*^95^ and *w*^1118^ control flies we should be able to characterize the impact of PNPO deficiency on VB6 metabolism. Furthermore, seizures in PNPO deficient patients cease with PLP or PN treatment [17, 20, 23, 24]. Our previous studies have also shown that PLP or PN supplementation can rescue the conditional lethality of *sgll*^95^ flies [31]. To further correlate the PLP level with seizures and lethality, we measured PLP in flies reared on the sugar-only diet as well as the sugar-only diet supplemented with PN.

**Figure 3.**
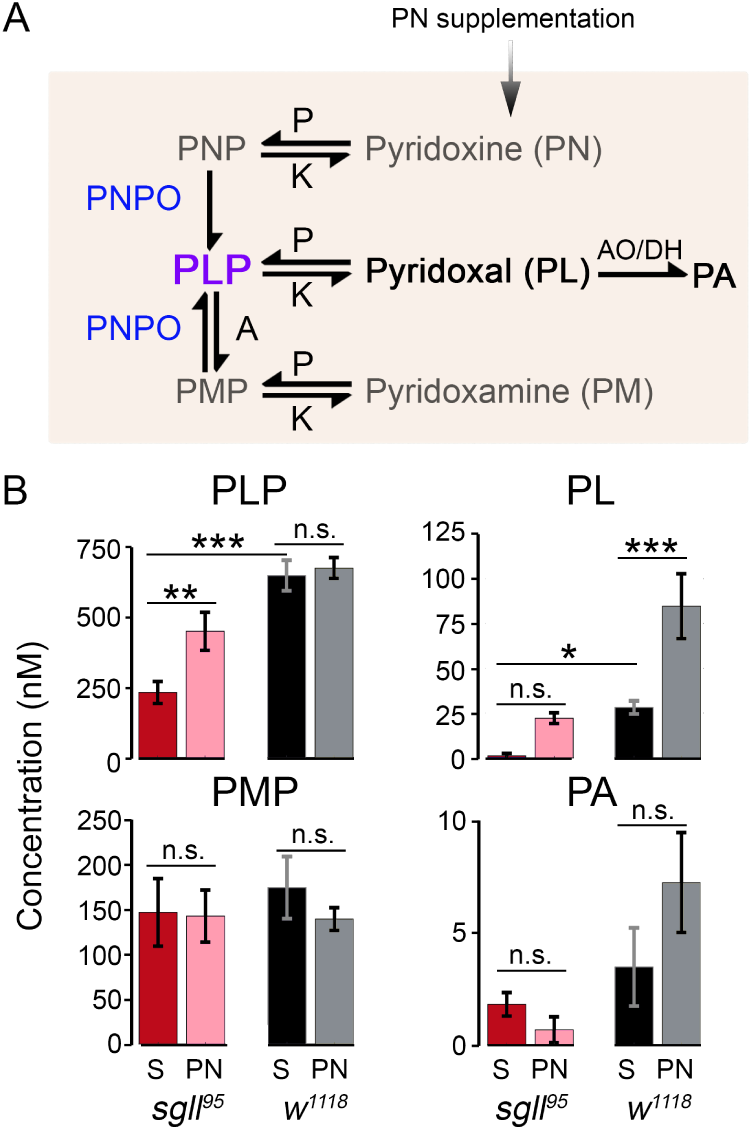
B6 vitamer measurements in *sgll*^95^ and *w*^1118^ flies under different conditions. A) A schematic representation of the interchangeable relationships among B6 vitamers from studies in mammals and humans. The conversion occurs primarily in the liver cells. After conversion, B6 vitamers (non-phosphorylated forms) are released into the circulation and transported to other tissues/organs. Among them, PL is the main form of VB6 for most tissues/organs. Excess PL is oxidized by aldehyde oxidase (AO) or NAD^+^-dependent aldehyde dehydrogenase (DH) to pyridoxic acid (PA) in the liver or kidney. PA is then excreted in urine. In normal humans, abundant B6 forms are PLP, PL, and PA. Sometimes PM or PMP are detectable, whereas PN or PNP usually are undetectable [49, 50, 51, 52, 53]. PNP: pyridoxine 5’-phosphate, PLP: pyridoxal 5’-phosphate, PMP: pyridoxamine 5’-phosphate, PNPO: pyridox(am)ine 5’-phosphate oxidase, P: phosphatase, K: pyridoxal kinase, A: aminotransferase. B) Six B6 vitamers (PN, PNP, PL, PLP, PM, and PMP) and PA were measured in whole flies. The relative B6 vitamer composition in *w*^1118^ flies on the sugar-only diet is PLP > PMP > PL > PA. The other B6 vitamers were undetectable. Levels of various B6 vitamers in *sgll*^95^ and *w*^1118^ flies under the sugar-only diet (S) or the sugar-only diet supplemented with PN (PN) are shown in four panels. ** *P* < 0.01, *** *P* < 0.001, n.s.: *P* > 0.05; Two-way ANOVA with Tukey’s *post hoc*, n = 3 biological replicates per genotype per condition.

As shown in Figure 3B, when reared on the sugar-only diet (S), *sgll*^95^ flies had significantly reduced PLP compared to *w*^1118^ flies (red bar vs. black bar, *P* = 4.020e-5), which was significantly improved by PN supplementation (red vs. pink bar, *P* = 0.0035). On the other hand, PN supplementation did not change the PLP level in *w*^1118^ flies (black vs. gray bar, *P* = 0.9126), suggesting that PLP is highly regulated in normal flies. The regulation of PLP in *w*^1118^ flies is most likely mediated through the conversion from PLP to PL as indicated by a significantly increased PL level after PN supplementation in *w*^1118^ flies compared to *sgll*^95^ flies (gene by treatment interaction: *P* = 0.0115; *P* = 0.0004 and *P* = 0.0988 the treatment effect in *w*^1118^ and *sgll*^95^ flies, respectively). In comparison, PMP and PA levels show no difference between *w*^1118^ and *sgll*^95^ flies before PN supplementation (*P* = 0.6914 and *P* = 0.5305 for PMP and PA, respectively). After PN supplementation, PA is significantly increased in *w*^1118^ flies compared to *sgll*^95^ flies (gene by treatment interaction: *P* = 0.0198; *P* = 0.0504 and *P* = 0.8017 for the treatment effect in *w*^1118^ and *sgll*^95^ flies, respectively), whereas PMP is similar between the two genotypes (gene by treatment interaction: *P* = 0.4062; *P* = 0.5159 and *P* = 0.9980 for the treatment effect in *w*^1118^ and *sgll*^95^ flies, respectively).

### PNPO is functionally conserved between humans and flies

Amino acid sequence alignment analysis revealed that the protein product of *sgll* shares more than 75% similarity with human PNPO [31]. To further study if the PNPO enzyme is functionally conserved between humans and flies, we generated transgenic flies by sub-cloning cDNAs from wild-type (WT) human and *Drosophila PNPO* gene into a pUAST vector (UAS-*hPNPO* and UAS-*sgll*, respectively). We used the GAL4/UAS system [54] to drive the expression of transgenes. To examine the effect of the ubiquitous expression of each transgene, we first backcrossed each UAS line and an *actin*-Gal4 line (genotype: *actin*-Gal4/*CyO*) to the *sgll*^95^ background. We then crossed each backcrossed UAS line with the backcrossed *actin*-Gal4 line to generate flies in which the transgene was ubiquitously expressed on the *sgll*^95^ background. The conditional survival rates of these flies (genotypes: *actin*-Gal4/UAS-*sgll*; *sgll*^95^/*sgll*^95^ flies and *actin*-Gal4/UAS-*hPNPO*; *sgll*^95^/*sgll*^95^) and control flies (genotypes: +/UAS-*hPNPO*; *sgll*^95^/*sgll*^95^, +/UAS-*sgll*; *sgll*^95^/*sgll*^95^, and *actin*-Gal4/+; *sgll*^95^/*sgll*^95^ and *sgll*^95^/*sgll*^95^) were examined.

In contrast to *sgll*^95^ homozygotes that died within five days on the sugar-only diet (Figure 4A, red curve), *sgll*^95^ flies with ubiquitous expression of WT *sgll* or *hPNPO* (i.e. rescue flies) displayed no mortality (blue or green circle versus red point-up triangle; *P* < 0.001 for both genotypes compared to *sgll*^95^ homozygotes) and did not show any seizure-like behavior. To confirm that, we performed behavioral characterization on rescue flies reared on the sugar-only diet for two or five days. As shown in Figure 4B-D, rescue flies behaved normally and had similar average speed correlation coefficients to *sgll*^95^ flies on the normal diet. The average speed correlation coefficient on Day 2 is 0.601 ±0.141 (Mean ± SD) and 0.702 ± 0.106 for *actin*-Gal4/UAS-*sgll*; *sgll*^95^/*sgll*^95^ (n = 14) and *actin*-Gal4/UAS-*hPNPO*; *sgll*^95^/*sgll*^95^ (n = 11), respectively. The average speed correlation coefficient on Day 5 is 0.722 ± 0.107 and 0.675 ± 0.160 for *actin*-Gal4/UAS-*sgll*; *sgll*^95^/*sgll*^95^ (n = 10) and *actin*-Gal4/UAS-*hPNPO*; *sgll*^95^/*sgll*^95^ (n = 10), respectively. These coefficients are not significantly different from those in *sgll*^95^ flies on the normal fly diet (*P* = 0.0766, *P* = 0.9866, *P* = 0.6873, and *P* = 0.6852, unpaired *t*-test). Furthermore, recordings of DLM activities from *actin*-Gal4/ UAS-*sgll*; *sgll*^95^/*sgll*^95^ and *actin*-Gal4/UAS-*hPNPO*; *sgll*^95^/*sgll*^95^ flies did not show the spiking burst discharges characteristic of *sgll*^95^ flies (Figure 4E, n = 7 for each genotype). Together, these data indicate that ubiquitous expression of *hPNPO* completely rescued the conditional seizures and lethality of *sgll*^95^ homozygotes and demonstrate the functional conservation of the PNPO enzyme in humans and flies.

**Figure 4.**
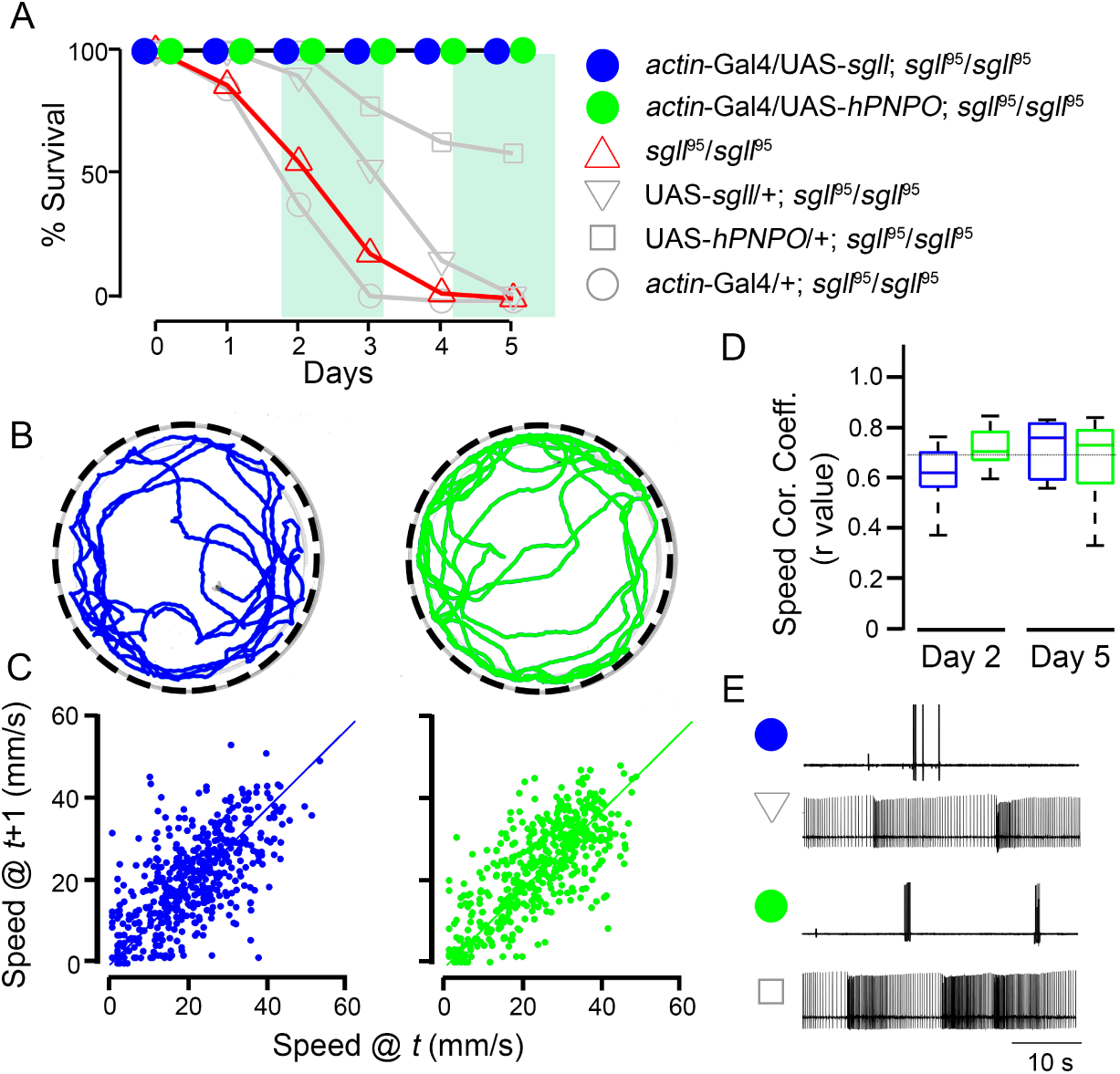
Ubiquitous expression of WT *sgll* or *hPNPO* completely rescued the lethality and seizures of *sgll*^95^ homozygotes. A) Survival of flies with various genotypes on the sugar-only diet. Ubiquitous expression of WT *sgll* or *hPNPO* completely rescued the conditional lethality of *sgll*^95^ homozygotes (blue or green circle vs. red point-up triangle, *P* < 0.0001, Log-rank test, n = 51-101). The curve of *sgll*^95^ homozygotes was replotted from Figure 1A. Similar to *sgll*^95^ homozygotes, *actin*-Gal4/+; *sgll*^95^/*sgll*^95^ (gray circle, n = 52) and UAS-*sgll*/+; *sgll*^95^/*sgll*^95^ (gray point-down triangle, n = 253) also died by day 5. Note that some UAS-*hPNPO*/+; *sgll*^95^/*sgll*^95^ flies survived by day 5 (gray square, total n = 314), which was presumably due to the expression of *hPNPO* in cells labeled by *Ddc*-Gal4 since the mutation in the *sgll*^95^ allele was identified in the *Ddc*-Gal4 driver line [31]. The green block indicates the time window in which the seizure characterizations were performed. Representative travel traces from *actin*-Gal4/UAS-*sgll*; *sgll*^95^/*sgll*^95^ (left) and *actin*-Gal4/UAS-*hPNPO*; *sgll*^95^/*sgll*^95^ (right). Speed correlation plots between the frame *t* and frame *t*+1. Each plot is corresponding to the one in panel B. D) Summary plot of speed correlation coefficient (Speed Cor. Coeff.) measured on day 2 or day 5. The black dotted line indicates the average speed correlation coefficient of *sgll*^95^ homozygotes on the normal diet (shown in Figure 1E). E) Representative spike traces of DLM spiking in rescue and their parental UAS lines.

### PNPO in the brain is necessary for normal brain function

The brain obtains VB6 from the circulation [12, 13]. Previous studies have shown that PLP and PL are the primary forms of VB6 in the circulation [55, 56], raising the question whether PNPO in the brain is necessary for normal brain functions.

To answer this question, we generated neural-specific *sgll* KD flies using an *elav*-Gal4 driver [57]. KD and corresponding control flies were then tested on the sugar-only diet. We observed both lethality (Figure 5A, *P* < 0.001) and seizures (Figure 5B) in *sgll* KD flies, both of which were absent from controls. However, compared to *sgll*^95^ or ubiquitous *sgll* KD flies (Figure 1A and Figure S1 A), neural-specific *sgll* KD flies were not as severely impaired. One possible explanation is that neural-specific *sgll* KD was not efficient. Since it was not feasible to quantify cell type-specific *sgll* KD efficiency precisely, we generated neural-specific *sgll* knockout (KO) flies using guide RNA stocks generated by the Transgenic RNAi project (https://fgr.hms.harvard.edu) and tested them on the sugar-only diet. As shown in Figure S2, neural-specific *sgll* KO flies also showed lethality. The survival rates of neural-specific *sgll* KD flies and neural-specific *sgll* KO flies are comparable, suggesting that the knockdown efficiency was high. It’s worth noting that in the ubiquitous *sgll* KD flies, the knockdown efficiency was over 90% as measured by the mRNA level [31]. Therefore, expression of PNPO in neurons is essential for both seizure prevention and survival but PNPO expression in other cell types also contributes. Since PLP can be converted to PL, cell membranes are permeable to PL, and PL can be converted to PLP independent of PNPO, it is conceivable that PNPO functions are only partially cell-autonomous.

**Figure 5.**
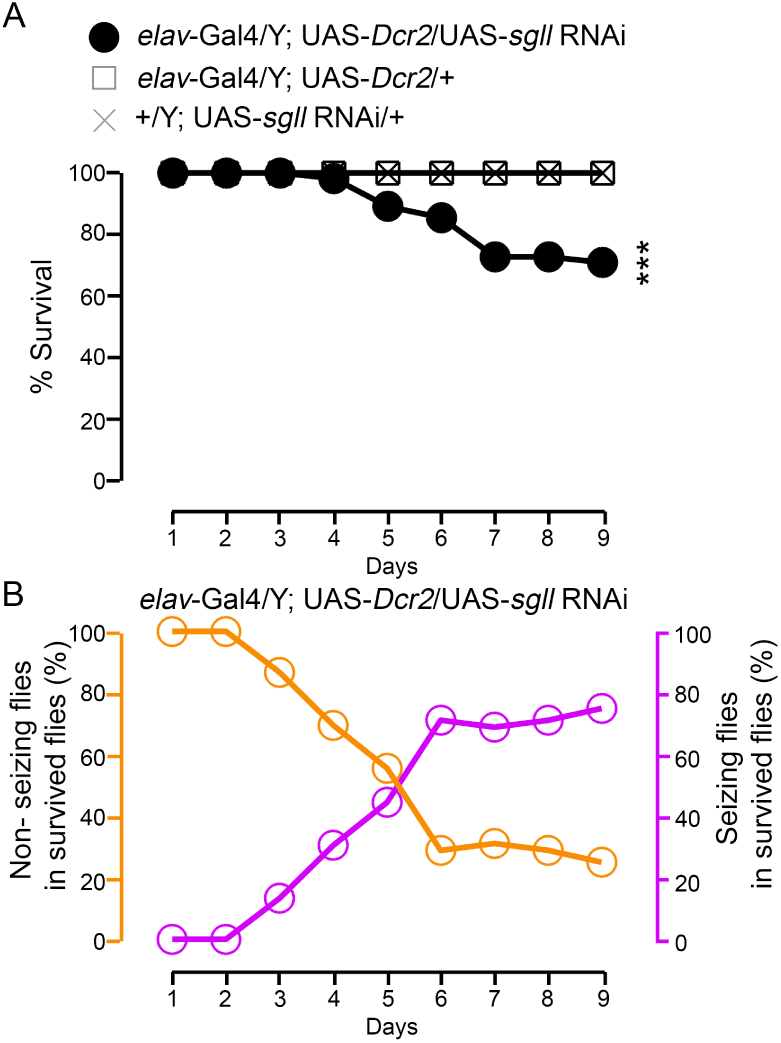
Survival and seizure rate of neural-specific *sgll* KD and control flies on the sugar-only diet. A) Survival of neural-specific *sgll* KD (genotype: *elav*-Gal4/Y;UAS-*Dcr2*/UAS-*sgll* RNAi) and control flies (genotypes: *elav*-Gal4/Y; UAS-*Dcr2*/+ and +/Y; UAS-*Dcr2*/UAS-*sgll* RNAi). *** *P* < 0.001, Log-rank test, compared to control lines, n = 55 - 60 per genotype. B) More than 70 % of survived neural-specific *sgll* KD flies showed seizure-like phenotype by day 9.

## Discussion

The identification of a hypomorphic *PNPO* allele, *sgll*^95^, in *Drosophila* by our earlier work [31] has opened up opportunities to study PNPO functions in an animal model for the first time. Here we report that PNPO deficiency causes seizures in *Drosophila*, similar to human conditions associated with PNPO deficiency. The seizure and lethality phenotypes are associated with low PLP levels and are completely rescued by VB6 supplementation and by ubiquitous expression of either WT *sgll* or WT human *PNPO* (*hPNPO*). Cell type-specific KD further suggests that PNPO in the brain is necessary for seizure prevention and survival.

The amino acid sequence of PNPO is evolutionarily conserved [31, 58]. The fact that PNPO deficiency induces seizures in flies as it does in humans (Figures 1 & 2) demonstrates that the biological function of PNPO is also conserved. Complete rescue of the conditional lethality and seizures of *sgll*^95^ flies by ubiquitous expression of *hPNPO* further confirmed this notion (Figure 4). All these provide a solid foundation for studying the fundamental biology of PNPO (Figure 5) and for future characterizations of human-disease-associated hPNPO mutations in *Drosophila* models.

NEE caused by severe PNPO deficiency is lethal if untreated. However, favorable outcomes are still possible if patients could be diagnosed and treated timely [59]. A sensitive and specific biomarker will undoubtedly help in the diagnosis. Given that PNPO determines the synthesis of PLP, the PLP level along with other B6 vitamers has been scrutinized in both PNPO deficient patients and individuals without PNPO deficiency. However, results have not been consistent for PLP level measurements [26], which can be attributed to the heterogeneous genetic background, different diets, as well as other uncontrolled factors. Recently, it has been shown that PNPO deficient patients have high plasma levels of PM [49], but the sample size is relatively small (n = 6) and majority of them (5 / 6) were on either PN or PN+PLP treatment. More samples are needed to validate this finding. In our fly models with well-controlled genetic background and environmental conditions, it is clear that PLP level is the best predictor of symptoms among B6 vitamers (Figure 3). It is worth noting that PMP instead of PM was detected in flies under the current conditions and the PMP level is not changed by PN supplementation and is not altered in *sgll*^95^ flies compared to *w*^1118^ controls.

Little is known about the neurobiological mechanisms underlying PNPO-deficiency-induced seizures. In humans, different metabolites have been measured in PNPO deficient patients, including those involved in the synthesis of dopamine, serotonin, GABA, and other amino acids such as threonine and glycine. A potential contributor to the seizure phenotype in PNPO deficiency flies is an altered inhibitory central circuit. Notably, our spike pattern analysis suggests that seizures in PNPO deficient flies (Figure 2) share characteristics with spike discharges in wild-type flies administered with picrotoxin, a GABA_*A*_ antagonist [47], supporting the notion of a potential role of dysfunctional GABAergic neurons as a consequence of PNPO deficiency. This is not surprising as PLP is required for GABA synthesis and GABA deficiency is known to be involved in the development of epilepsy.

This interpretation is also in agreement with results from animal models of other gene mutations in the VB6 pathway. Other than PNPO, several genes have been identified in VB6-dependent epilepsy patients, including *ALDH7A1* [60], *ALPL* [61], and *PROSC* [62, 63], which encodes aldehyde dehydrogenase 7A1, tissue non-specific alkaline phosphatase (TNSALP), and proline synthetase co-transcribed homolog, respectively. These gene products affect the availability of PLP through different mechanisms. Knockout animal models have been generated to study the function of *ALPL* [64] and *ALDH7A1* genes [65]. Those studies suggest that reduced intracellular PLP and in consequence, the GABA deficiency are likely the main contributors to their seizure phenotype.

Seizures have long been studied in *Drosophila* [32]. Fly models based on mutations in a number of genes have been established and well characterized, including genes encoding sodium, potassium, and calcium channels [66, 67, 39], as well as genes encoding proteins that can affect the functions of ion transporters [68, 69, 70]. The seizure activity *in sgll* is distinct from two major classes of hyperexcitable *Drosophila* mutants, bang-sensitive mutants (e.g. *bss, eas* [33]) and temperature-sensitive mutants (e.g. sei [38], *cac* [39, 40], *Shu* [41]), in which wing-buzzing and DLM spiking can be triggered by mechanical shock or high temperature exposure. In sgll mutants, wing-buzzing and DLM spike discharges appear spontaneously. Notably, the spontaneous DLM spike discharges in sgll mutants are reminiscent to spike patterns recruited by injection of the GABA_*A*_ antagonist picrotoxin in WT flies [47] (Figure 2C). However, additional work is required to demonstrate the role of GABAergic circuitry in driving seizure discharges in *sgll* mutants.

Characterization of PNPO mutants makes it possible to examine genetic/functional interactions between PNPO and other proteins implicated in epileptogenesis. For example, VB6 has been shown to control seizures in human patients who carry mutations in *KCNQ2* [71, 72] or CACNA1A [73], both of which encode voltage-gated ion channels. Detailed analysis of the VB6 metabolism pathway in known epilepsy models could reveal a better understanding of how seizures arise in these seizure-prone patients and potentially provide new therapeutic avenues to suppress seizure activity.

Epilepsy affects more than 70 million people in the world with one-third of them not well-controlled by drug treatments [74, 75]. Now with the advent of next-generation sequencing, treatments may be designed based on genetic information. The power of such an approach is especially exciting for defects in genes encoding metabolic enzymes as treatments may include simple dietary changes. Thus, valid animal models are valuable tools for testing treatment options, and help to elucidate the fundamental biology of these causal genes.

## Methods

### *Drosophila* strains

Electrophysiology experiments were performed on flies bred on Frankel & Brosseau’s media [76] at the University of Iowa. For all other experiments, flies were bred on the standard cornmeal-yeast-molasses medium from Fly Kitchen at the University of Chicago. Flies used in all experiments were raised and tested at room temperature (∼23 *°*C).The *sgll*^95^ line and its genetic control wild-type line, *w*^1118^, were described before [31]. The UAS-*sgll* RNAi transgenic line (VDRC #105941), as well as its control, was obtained from the Vienna Drosophila Resource Center (VDRC). The *actin*-Gal4 (BDSC #4414), *elav*-Gal4 (#25750), *elav*-Gal4; UAS-*Cas9*/*CyO* (#67073), and *sgll* gRNA (#79746) were obtained from the Bloomington Drosophila Stock Center (BDSC) at Indiana.

### Generation of transgenic flies

To generate transgenic flies, we constructed UAS-*sgll* and UAS-*hPNPO* by sub-cloning the cDNAs of *Drosophila* wild-type *sgll* and human wild-type *PNPO*, respectively, into pUAST and injected into *w*^1118^ commercially (Rainbow Transgenic Flies, Inc). Specifically, fly WT *sgll* cDNA (RA form, see reference [31]) were amplified from *w*^1118^ cDNAs with primers: 5’-CGA ATT CGC CAC CAT GAA GTT GCT GCA AAC AAT TCG AAG G-3’ and 5’-AGA GCT CCT AAG GAG CCA GCC GTT CGT ACA CCC A-3’; and human WT *hPNPO* cDNA was amplified from human brain cDNA library (TaKaRa, Cat #637242) with primers: 5’-CGA ATT CGC CAC CAT GAC GTG CTG GCT GCG GGG CGT CA-3’ and 5’-GGA GCT CTT AAG GTG CAA GTC TCT CAT AGA GCC AGT CTT-3’. The sequences of constructs were confirmed by Sanger sequencing at the DNA Sequencing Facility at the University of Chicago.

### Survival study

We followed our previous method [31]. In brief, male flies, 1-3-day-old, were anesthetized briefly and transferred into a culture vial filled with 4% sucrose in 1% agar. Fifteen to twenty flies were grouped into a vial. Survival was checked daily.

### Behavioral recording and data analysis

For recordings in *sgll*^95^ or rescue flies, 1-2-day-old male flies were anesthetized briefly and maintained on 4% sucrose in 1% agar for two or five days as indicated in the text. For recordings in ubiquitous *sgll* KD flies, we used female flies because male flies died sooner and gave a shorter time window to do recordings (Figure S1). Therefore, 1-2-day-old female *sgll* KD flies were anesthetized briefly and maintained on 4% sucrose in 1% agar for two days. After treatment, each individual fly was transferred without anesthetization to a 60-mm petri dish (Corning, Cat #430166) for recording. The dish was pre-filled with 1% agar to maintain humidity. Each fly was recorded for 3 min at 20 frames per second using a Flea3 video camera (FLIR Integrated Imaging Solutions, Inc). Videos were saved to the computer with FlyCapture (FLIR Integrated Imaging Solutions, Inc) and tracked with the IowaFLI Tracker [42]. After tracking, the travel traces and scatter plots were plotted in Matlab (R2018b, Mathworks, Natick, MA). For clarity, only the first five hundred frames were plotted in the scatter plots shown in Figures 1, Figure S1, and Figure 4.

### Electrophysiological recording of DLM flight muscle activity

We followed the protocol from our previous publications [43, 46]. In brief, a single fly, either male or female, was shortly anesthetized on ice and glued to a tungsten wire between neck and thorax. After a 30-minute recovery period, two sharpened tungsten electrodes were inserted into the left and right thorax, one on each side, targeting the top-most dorsal longitudinal muscle (DLMa, [77]). A reference electrode was inserted into the abdomen of the fly. Electrical activity in each muscle were picked up with an AC amplifier (AM Systems Model 1800, Carlsbourg, WA) and digitized by data acquisition card (USB 6210 National Instruments, Austin TX) controlled by LabVIEW8.6 (National Instruments). Spike detection was done using a custom-written Matlab script. Following the conventions in our previous publication [47], the instantaneous firing frequency for an individual spike was defined as the reciprocal of the inter-spike interval between the current spike and the succeeding spike (ISI^−1^). The instantaneous coefficient of variation, CV_2_ [78] for a pair of ISI^−1^ corresponding to adjacent interval i and I +1 was shown as:

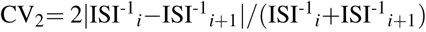

Lower CV_2_ values indicate ISI^−1^ values with little variability, and higher CV_2_ values correspond to irregular ISI^−1^ values [47].

### B6 vitamer measurements

For measuring levels of B6 vitamers, a group of 20 male *sgll*^95^ or *w*^1118^ flies, 1-3-day-old, were picked into vials supplied with 4% sucrose in 1% agar, which was supplemented with or without 500 ng/ml of PN. The concentration of 500 ng/ml was selected based on our previous studies ([31]) showing that all flies survive under this condition. Forty-eight hours later, whole flies in each vial were homogenized in cold trichloroacetic acid. After centrifugation, the supernatant in each vial was collected into new Eppendorf tubes, which were frozen at −80 *°*C until B6 vitamer measurements. The concentration of each B6 vitamer in fly lysates was quantified by Ultra-Performance-Liquid-Chromatography-Tandem-Mass-Spectrometry (UPLC-MS/MS) [79].

### Survival and seizures in neural-specific *sgll* KD and control flies

For survival, 1-3-day-old male flies, were anesthetized briefly and transferred into a culture vial filled with 4% sucrose in 1% agar. Fifteen to twenty flies were grouped into a vial. Survival was checked daily. We also recorded the number of flies that showed seizure phenotypes during a 2-minute-time window each day. Seizure phenotypes include wing-buzzing, body-rolling, and wings-up, all of which were described in Figure 1.

### Statistical analysis

Statistical analysis was performed using R (version 3.3.2). Details on statistical analyses, including sample sizes, tests performed, exact *P* values, and multiple tests correction if necessary, are provided within the figure legends or in the text describing each figure.

## Acknowledgments

We thank Dr. Edwin Ferguson (Department of Molecular Genetics and Cell Biology) for commenting on the manuscript. We thank Dr. Wei Du (Ben May Department for Cancer Research) for providing pUAST plasmids. We thank Qian Yang, Susie A. Turkson, Chris Wreden, Hanyu Li, Wenhan Chang, Wei Liu, and Wenqin Fu for technical support. We thank the Bloomington Drosophila Stock Center and the Vienna Drosophila Resource Center for *Drosophila* stocks. We thank Rainbow Transgenic Flies, Inc. for generating transgenic flies. This work was supported by the National Institutes of Health [T32MH020065 to W.C., R01GM100768 to X.Z., AG051513 to C.F.W]; the Donald F. Steiner Scholarship Fund (to W.C.); and Iowa Neuroscience Institute Fellowship (to A.I.).

## Author contributions

W.C. and X.Z. conceived the project. W.C. wrote the first draft of the manuscript. All authors edited the manuscript. W.C. designed and conducted rescue experiments, behavioral characterization of seizures, and knockdown experiments. C.-F.W. and A.I. designed the electrophysiological studies. A.I. and W.C. conducted electrophysiological characterization of seizures and A.I. performed data analysis. M.A., M.B., and N.V.D. conducted VB6 measurements. W.C. and A.I. constructed figures and performed statistical analyses.

## References

[1] Philippa B Mills, Robert AH Surtees, Michael P Champion, Clare E Beesley, Neil Dalton, Peter J Scambler, Simon JR Heales, Anthony Briddon, Irene Scheimberg, Georg F Hoffmann, et al. Neonatal epileptic encephalopathy caused by mutations in the pnpo gene encoding pyri-dox (am) ine 5-phosphate oxidase. Human molecular genetics, 14(8):1077–1086, 2005.

[2] Jacques L Michaud, Mathieu Lachance, Fadi F Hamdan, Lionel Carmant, Anne Lortie, Paola Diadori, Philippe Major, Inge A Meijer, Emmanuelle Lemyre, Patrick Cossette, et al. The genetic landscape of infantile spasms. Human molecular genetics, 23(18):4846–4858, 2014.

[3] Philippa B Mills, Stephane SM Camuzeaux, Emma J Footitt, Kevin A Mills, Paul Gissen, Laura Fisher, Krishna B Das, Sophia M Varadkar, Sameer Zuberi, Robert McWilliam, et al. Epilepsy due to pnpo mutations: geno-type, environment and treatment affect presentation and outcome. Brain, 137(5):1350–1360, 2014.

[4] Martino L di Salvo, Mario Mastrangelo, Isabel Nogués, Manuela Tolve, Alessandro Paiardini, Carla Carducci, Davide Mei, Martino Montomoli, Angela Tramonti, Renzo Guerrini, et al. Pyridoxine-5-phosphate oxidase (pnpo) deficiency: Clinical and biochemical alterations associated with the c. 347g> a (p.*·* arg116gln) mutation. Molecular genetics and metabolism, 122(1-2):135–142, 2017.

[5] Jiao Xue, Xingzhi Chang, Yuehua Zhang, and Zhixian Yang. Novel phenotypes of pyridox (am) ine-5’-phosphate oxidase deficiency and high prevalence of c. 445 448del mutation in chinese patients. Metabolic brain disease, 32(4):1081–1087, 2017.

[6] Esmond E Snell, Beverly M Guirard, Roger J Williams, et al. Occurrence in natural products of a physiologically active metabolite of pyridoxine. Journal of Biological Chemistry, 143:519–530, 1942.

[7] Stanton A Harris, Dorothea Heyl, Karl Folkers, et al. The structure and synthesis of pyridoxamine and pyridoxal. Journal of Biological Chemistry, 154:314–316, 1944.

[8] Riccardo Percudani and Alessio Peracchi. A genomic overview of pyridoxal-phosphate-dependent enzymes. EMBO reports, 4(9):850–854, 2003.

[9] Marcelina Parra, Seth Stahl, and Hanjo Hellmann. Vitamin b6 and its role in cell metabolism and physiology. Cells, 7(7):84, 2018.

[10] Jeong Han Kang, Mi-Lim Hong, Dae Won Kim, Jin-seu Park, Tae-Cheon Kang, Moo Ho Won, Nam-In Baek, Byung Jo Moon, Soo Young Choi, and Oh-Shin Kwon. Genomic organization, tissue distribution and deletion mutation of human pyridoxine 5-phosphate oxidase. European journal of biochemistry, 271(12):2452–2461, 2004.

[11] Alfred H Merrill and J Michael Henderson. Vitamin b6 metabolism by human livera. Annals of the New York Academy of Sciences, 585(1):110–117, 1990.

[12] Reynold Spector. Vitamin b6 transport in the central nervous system: in vitro studies. Journal of neurochemistry, 30(4):889–897, 1978.

[13] Reynold Spector. Vitamin b6 transport in the central nervous system: in vivo studies. Journal of neurochemistry, 30(4):881–887, 1978.

[14] G. F. Hoffmann, B. Schmitt, M. Windfuhr, N. Wagner, H. Strehl, S. Bagci, A. R. Franz, P. B. Mills, P. T. Clayton, M. R. Baumgartner, B. Steinmann, T. Bast, N. I. Wolf, and Johannes Zschocke. Pyridoxal 5-phosphate may be curative in earlyonset epileptic encephalopathy. J. Inherit. Metab. Dis., 30(1):96–99, 2007.

[15] Angeles Ruiz, Judit García-Villoria, Aida Ormazabal, Johannes Zschocke, Miquel Fiol, Aleix Navarro-Sastre, Rafael Artuch, Ma Antonia Vilaseca, and Antonia Ribes. A new fatal case of pyridox (am) ine 5-phosphate oxidase (pnpo) deficiency. Molecular genetics and metabolism, 93(2):216–218, 2008.

[16] Morad Khayat, Stanley H. Korman, Pnina Frankel, Zal-man Weintraub, Sylvia Hershckowitz, Vered Fleisher Sheffer, Mordechai Ben Elisha, Ronald A. Wevers, and Tzipora C. Falik-Zaccai. PNPO deficiency: An under diagnosed inborn error of pyridoxine metabolism. Mol. Genet. Metab., 94(4):431–434, 2008.

[17] Phillip L Pearl, Keith Hyland, J Chiles, Colleen L McGavin, Yuezhou Yu, and Donald Taylor. Partial pyridoxine responsiveness in pnpo deficiency. In JIMD Reports–Case and Research Reports, 2012/6, pages 139–142. Springer, 2012.

[18] Stephanie Porri, Joel Fluss, Barbara Plecko, Eduard Paschke, Christian M Korff, and Ilse Kern. Positive outcome following early diagnosis and treatment of pyridoxal-5-phosphate oxidase deficiency: a case report. Neuropediatrics, 45(01):064–068, 2014.

[19] M Veeravigrom, P Damrongphol, R Ittiwut, C Ittiwut, K Suphapeetiporn, and V Shotelersuk. Pyridoxal 5-phosphate-responsive epilepsy with novel mutations in the pnpo gene: a case report. Genetics and Molecular Research, 14(4):14130–14135, 2015.

[20] Barbara Plecko, Karl Paul, Philippa Mills, Peter Clayton, Eduard Paschke, Oliver Maier, Oswald Hasselmann, Gudrun Schmiedel, Simone Kanz, Mary Connolly, et al. Pyridoxine responsiveness in novel mutations of the pnpo gene. Neurology, 82(16):1425–1433, 2014.

[21] Bernhard Schmitt, Matthias Baumgartner, Philippa B Mills, Peter T Clayton, Cornelis Jakobs, Elmar Keller, and Gabriele Wohlrab. Seizures and paroxysmal events: symptoms pointing to the diagnosis of pyridoxine-dependent epilepsy and pyridoxine phosphate oxidase deficiency. Developmental Medicine & Child Neurology, 52(7):e133–e142, 2010.

[22] S Bagci, J Zschocke, G F Hoffmann, T Bast, J Klepper, A Müller, A Heep, P Bartmann, and a R Franz. Pyridoxal phosphate-dependent neonatal epileptic encephalopathy. Arch. Dis. Child. Fetal Neonatal Ed., 93(2):F151–2, 2008.

[23] Tyson L Ware, John Earl, Gajja S Salomons, Eduard A Struys, Heidi L Peters, Katherine B Howell, James J Pitt, and Jeremy L Freeman. Typical and atypical phenotypes of pnpo deficiency with elevated csf and plasma pyridoxamine on treatment. Developmental Medicine & Child Neurology, 56(5):498–502, 2014.

[24] Andrea Guerin, Aly S Aziz, Carly Mutch, Jillian Lewis, Cristina Y Go, and Saadet Mercimek-Mahmutoglu. Pyridox (am) ine-5-phosphate oxidase deficiency treatable cause of neonatal epileptic encephalopathy with burst suppression: case report and review of the literature. Journal of child neurology, 30(9):1218–1225, 2015.

[25] Monisha Goyal, Pierre R Fequiere, Tony M McGrath, and Keith Hyland. Seizures with decreased levels ofpyridoxal phosphate in cerebrospinal fluid. Pediatric neurology, 48(3):227–231, 2013.

[26] Alina Levtova, Stephane Camuzeaux, Anne-Marie Laberge, Pierre Allard, Catherine Brunel-Guitton, Paola Diadori, Elsa Rossignol, Keith Hyland, Peter T Clayton, Philippa B Mills, et al. Normal cerebrospinal fluid pyridoxal 5-phosphate level in a pnpo-deficient patient with neonatal-onset epileptic encephalopathy. In JIMD Reports, Volume 22, pages 67–75. Springer, 2015.

[27] B Jaeger, NG Abeling, GS Salomons, EA Struys, M Simas-Mendes, VG Geukers, and BT Poll-The. Pyridoxine responsive epilepsy caused by a novel homozygous pnpo mutation. Molecular genetics and metabolism reports, 6:60–63, 2016.

[28] Martino L di Salvo, Mario Mastrangelo, Isabel Nogués, Manuela Tolve, Alessandro Paiardini, Carla Carducci, Davide Mei, Martino Montomoli, Angela Tramonti, Renzo Guerrini, et al. Biochemical data from the characterization of a new pathogenic mutation of human pyridoxine-5’-phosphate oxidase (pnpo). Data in brief, 15:868–875, 2017.

[29] Annapurna Sudarsanam, Harry Singh, Bridget Wilcken, Michael Stormon, Susan Arbuckle, Bernhard Schmitt, Peter Clayton, John Earl, and Richard Webster. Cirrhosis associated with pyridoxal 5-phosphate treatment of pyridoxamine 5-phosphate oxidase deficiency. In JIMD Reports, Volume 17, pages 67–70. Springer, 2014.

[30] D Coman, P Lewindon, P Clayton, and K Riney. Pnpo deficiency and cirrhosis: expanding the clinical phenotype? In JIMD Reports, Volume 25, pages 71–75. Springer, 2015.

[31] Wanhao Chi, Li Zhang, Wei Du, and Xiaoxi Zhuang. A nutritional conditional lethal mutant due to pyridoxine 5-phosphate oxidase deficiency in drosophila melanogaster. G3: Genes, Genomes, Genetics, 4(6):1147–1154, 2014.

[32] Seymour Benzer. From the gene to behavior. Jama, 218(7):1015–1022, 1971.

[33] Barry Ganetzky and Chun-Fang Wu. Indirect suppression involving behavioral mutants with altered nerve excitability in drosophila melanogaster. Genetics, 100(4):597–614, 1982.

[34] Paul Pavlidis, Mani Ramaswami, and Mark A Tanouye. The drosophila easily shocked gene: a mutation in a phospholipid synthetic pathway causes seizure, neuronal failure, and paralysis. Cell, 79(1):23–33, 1994.

[35] HaiGuang Zhang, Jeff Tan, Elaine Reynolds, Daniel Kuebler, Sally Faulhaber, and Mark Tanouye. The drosophila slamdance gene: a mutation in an aminopeptidase can cause seizure, paralysis and neuronal failure. Genetics, 162(3):1283–1299, 2002.

[36] Thomas A Grigliatti, Linda Hall, Raja Rosenbluth, and David T Suzuki. Temperature-sensitive mutations in drosophila melanogaster. Molecular and General Genetics MGG, 120(2):107–114, 1973.

[37] Martin G Burg and Chun-Fang Wu. Mechanical and temperature stressor–induced seizure-and-paralysis behaviors in drosophila bang-sensitive mutants. Journal of neurogenetics, 26(2):189–197, 2012.

[38] F Rob Jackson, Susan D Wilson, Gary R Strichartz, and Linda M Hall. Two types of mutants affecting voltage-sensitive sodium channels in drosophila melanogaster. Nature, 308(5955):189, 1984.

[39] Fumiko Kawasaki, Ryan Felling, and Richard W Ordway. A temperature-sensitive paralytic mutant defines a primary synaptic calcium channel in drosophila. Journal of Neuroscience, 20(13):4885–4889, 2000.

[40] Gabrielle E Rieckhof, Motojiro Yoshihara, Zhuo Guan, and J Troy Littleton. Presynaptic n-type calcium channels regulate synaptic growth. Journal of Biological Chemistry, 278(42):41099–41108, 2003.

[41] Garrett A Kaas, Junko Kasuya, Patrick Lansdon, Atsushi Ueda, Atulya Iyengar, Chun-Fang Wu, and Toshihiro Kitamoto. Lithium-responsive seizure-like hyperexcitability is caused by a mutation in the drosophila voltage-gated sodium channel gene paralytic. eNeuro, 3(3), 2016.

[42] Atulya Iyengar, Jordan Imoehl, Atsushi Ueda, Jeffery Nirschl, and Chun-Fang Wu. Automated quantification of locomotion, social interaction, and mate preference in drosophila mutants. Journal of neurogenetics, 26(3-4):306–316, 2012.

[43] Atulya Iyengar and Chun-Fang Wu. Flight and seizure motor patterns in drosophila mutants: simultaneous acoustic and electrophysiological recordings of wing beats and flight muscle activity. Journal of neurogenetics, 28(3-4):316–328, 2014.

[44] Michael H Dickinson and Michael S Tu. The function of dipteran flight muscle. Comparative Biochemistry and Physiology Part A: Physiology, 116(3):223–238, 1997.

[45] P Pavlidis and MA Tanouye. Seizures and failures in the giant fiber pathway of drosophila bang-sensitive paralytic mutants. Journal of Neuroscience, 15(8):5810–5819, 1995.

[46] Jisue Lee and Chun-Fang Wu. Electroconvulsive seizure behavior in drosophila: analysis of the physiological repertoire underlying a stereotyped action pattern in bang-sensitive mutants. Journal of Neuroscience, 22(24):11065–11079, 2002.

[47] Jisue Lee, Atulya Iyengar, and Chun-Fang Wu. Distinctions among electroconvulsion-and proconvulsant-induced seizure discharges and native motor patterns during flight and grooming: Quantitative spike pattern analysis in drosophila flight muscles. bioRxiv doi:https://doi.org/10.1101/481234, 2019.

[48] Réjean M Guerriero, Archana A Patel, Brian Walsh, Fiona M Baumer, Ankoor S Shah, Jurriaan M Peters, Lance H Rodan, Pankaj B Agrawal, Phillip L Pearl, and Masanori Takeoka. Systemic manifestations in pyridox (am) ine 5-phosphate oxidase deficiency. Pediatric neurology, 76:47–53, 2017.

[49] Déborah Mathis, Lucia Abela, Monique Albersen, Céline Bürer, Lisa Crowther, Karin Beese, Hans Hartmann, Lev-inus A Bok, Eduard Struys, Sorina M Papuc, et al. The value of plasma vitamin b6 profiles in early onset epileptic encephalopathies. Journal of Inherited Metabolic Disease: Official Journal of the Society for the Study of Inborn Errors of Metabolism, 39(5):733–741, 2016.

[50] Monique Albersen, Floris Groenendaal, Maria van der Ham, Tom J de Koning, Marjolein Bosma, Wouter F Visser, Gepke Visser, Nanda M Verhoeven-Duif, et al. Vitamin b6 vitamer concentrations in cerebrospinal fluid differ between preterm and term newborn infants. Pediatrics, 130(1):e191–e198, 2012.

[51] Emma J Footitt, Peter T Clayton, Kevin Mills, Simon J Heales, Viruna Neergheen, Marcus Oppenheim, and Philippa B Mills. Measurement of plasma b6 vitamer profiles in children with inborn errors of vitamin b6 metabolism using an lc-ms/ms method. Journal of Inherited Metabolic Disease: Official Journal of the Society for the Study of Inborn Errors of Metabolism, 36(1):139–145, 2013.

[52] Monique Albersen, Marjolein Bosma, Jurjen J Luykx, Judith JM Jans, Steven C Bakker, Eric Strengman, Paul J Borgdorff, Peter JM Keijzers, Eric Pa van Dongen, Peter Bruins, et al. Vitamin b-6 vitamers in human plasma and cerebrospinal fluid. The American journal of clinical nutrition, 100(2):587–592, 2014.

[53] Monique Albersen, Marjolein Bosma, Judith JM Jans, Floris C Hofstede, Peter M Van Hasselt, Gepke Visser, Nanda M Verhoeven-Duif, et al. Vitamin b6 in plasma and cerebrospinal fluid of children. PloS one, 10(3):e0120972, 2015.

[54] Andrea H Brand and Norbert Perrimon. Targeted gene expression as a means of altering cell fates and generating dominant phenotypes. development, 118(2):401–415, 1993.

[55] Walter B Dempsey and Halvor N Christensen. The specific binding of pyridoxal 5’-phosphate to bovine plasma albumin. Journal of Biological Chemistry, 237(4):1113–1120, 1962.

[56] Haile Mehansho and LM Henderson. Transport and accumulation of pyridoxine and pyridoxal by erythrocytes. Journal of Biological Chemistry, 255(24):11901–11907, 1980.

[57] David M Lin and Corey S Goodman. Ectopic and increased expression of fasciclin ii alters motoneuron growth cone guidance. Neuron, 13(3):507–523, 1994.

[58] Faik N Musayev, Martino L Di Salvo, Tzu-Ping Ko, Verne Schirch, and Martin K Safo. Structure and properties of recombinant human pyridoxine 5-phosphate oxidase. Protein Science, 12(7):1455–1463, 2003.

[59] J Hatch, D Coman, P Clayton, P Mills, S Calvert, RI Webster, and Kate Riney. Normal neurodevelopmental outcomes in pnpo deficiency: a case series and literature review. In JIMD Reports, Volume 26, pages 91–97. Springer, 2015.

[60] Philippa B Mills, Eduard Struys, Cornelis Jakobs, Barbara Plecko, Peter Baxter, Matthias Baumgartner, Michél AAP Willemsen, Heymut Omran, Uta Tacke, Birgit Uhlenberg, et al. Mutations in antiquitin in individuals with pyridoxine-dependent seizures. Nature medicine, 12(3):307, 2006.

[61] Maki Oyachi, Daisuke Harada, Natsuko Sakamoto, Kaoru Ueyama, Kawai Kondo, Kanako Kishimoto, Masafumi Izui, Yuiko Nagamatsu, Hiroko Kashiwagi, Miho Yamamuro, et al. A case of perinatal hypophosphatasia with a novel mutation in the alpl gene: clinical course and review of the literature. Clinical Pediatric Endocrinology, 27(3):179–186, 2018.

[62] Niklas Darin, Emma Reid, Laurence Prunetti, Lena Samuelsson, Ralf A Husain, Matthew Wilson, Basma El Yacoubi, Emma Footitt, WK Chong, Louise C Wilson, et al. Mutations in prosc disrupt cellular pyridoxal phosphate homeostasis and cause vitamin-b6-dependent epilepsy. The American Journal of Human Genetics, 99(6):1325–1337, 2016.

[63] Barbara Plecko, Markus Zweier, Anaïs Begemann, Deborah Mathis, Bernhard Schmitt, Pasquale Striano, Martina Baethmann, Maria Stella Vari, Francesca Beccaria, Federico Zara, et al. Confirmation of mutations in prosc as a novel cause of vitamin b 6-dependent epilepsy. Journal of medical genetics, 54(12):809–814, 2017.

[64] Katrina G Waymire, J Dennis Mahuren, J Michael Jaje, Tomás R Guilarte, Stephen P Coburn, and Grant R Mac-Gregor. Mice lacking tissue non–specific alkaline phosphatase die from seizures due to defective metabolism of vitamin b–6. Nature genetics, 11(1):45, 1995.

[65] Izabella A Pena, Yann Roussel, Kate Daniel, Kevin Mongeon, Devon Johnstone, Hellen Weinschutz Mendes, Marjolein Bosma, Vishal Saxena, Nathalie Lepage, Pranesh Chakraborty, et al. Pyridoxine-dependent epilepsy in zebrafish caused by aldh7a1 deficiency. Genetics, 207(4):1501–1518, 2017.

[66] Kate Loughney, Robert Kreber, and Barry Ganetzky. Molecular analysis of the para locus, a sodium channel gene in drosophila. Cell, 58(6):1143–1154, 1989.

[67] Steven A Titus, Jeffrey W Warmke, and Barry Ganetzky. The drosophila erg k+ channel polypeptide is encoded by the seizure locus. Journal of Neuroscience, 17(3):875–881, 1997.

[68] Michael J Palladino, Jill E Bower, Robert Kreber, and Barry Ganetzky. Neural dysfunction and neurodegener-ation indrosophila na+/k+ atpase alpha subunit mutants. Journal of Neuroscience, 23(4):1276–1286, 2003.

[69] Daria S Hekmat-Scafe, Miriam Y Lundy, Rakhee Ranga, and Mark A Tanouye. Mutations in the k+/clcotrans-porter gene kazachoc (kcc) increase seizure susceptibility in drosophila. Journal of Neuroscience, 26(35):8943–8954, 2006.

[70] Jan E Melom and J Troy Littleton. Mutation of a nckx eliminates glial microdomain calcium oscillations and enhances seizure susceptibility. Journal of Neuroscience, 33(3):1169–1178, 2013.

[71] Emma S Reid, Hywel Williams, Polona Le Quesne Stabej, Chela James, Louise Ocaka, Chiara Bacchelli, Emma J Footitt, Stewart Boyd, Maureen A Cleary, Philippa B Mills, et al. Seizures due to a kcnq2 mutation: treatment with vitamin b 6. In JIMD Reports, Volume 27, pages 79–84. Springer, 2015.

[72] Kerstin Alexandra Klotz, Johannes R Lemke, Rudolf Korinthenberg, and Julia Jacobs. Vitamin b6–responsive epilepsy due to a novel kcnq2 mutation. Neuropediatrics, 48(03):199–204, 2017.

[73] Xiaoping Du, You Chen, Yongxiong Zhao, Wei Luo, Zhidong Cen, and Weicheng Hao. Dramatic response to pyridoxine in a girl with absence epilepsy with ataxia caused by a de novo cacna1a mutation. Seizure-European Journal of Epilepsy, 45:189–191, 2017.

[74] Anuradha Singh and Stephen Trevick. The epidemiology of global epilepsy. Neurologic clinics, 34(4):837–847, 2016.

[75] Zhibin Chen, Martin J Brodie, Danny Liew, and Patrick Kwan. Treatment outcomes in patients with newly diagnosed epilepsy treated with established and new antiepileptic drugs: a 30-year longitudinal cohort study. JAMA neurology, 75(3):279–286, 2018.

[76] AWK Frankel and GE Brousseau. Drosophila medium that does not require dried yeast. Drosophila Information Service, 43:184, 1968.

[77] Jon D Levine and Malcolm Hughes. Stereotaxic map of the muscle fibers in the indirect flight muscles of drosophila melanogaster. Journal of morphology, 140(2):153–158, 1973.

[78] Gary R Holt, William R Softky, Christof Koch, and Rodney J Douglas. Comparison of discharge variability in vitro and in vivo in cat visual cortex neurons. Journal of neurophysiology, 75(5):1806–1814, 1996.

[79] M Van der Ham, M Albersen, TJ De Koning, G Visser, A Middendorp, M Bosma, NM Verhoeven-Duif, and MGM de Sain-van der Velden. Quantification of vitamin b6 vitamers in human cerebrospinal fluid by ultra performance liquid chromatography–tandem mass spectrometry. Analytica chimica acta, 712:108–114, 2012.

