## Supplementary material for "Pyridox(am)ine 5’-phosphate oxidase deficiency induces seizures in *Drosophila melanogaster*": Supplmental Figure 1

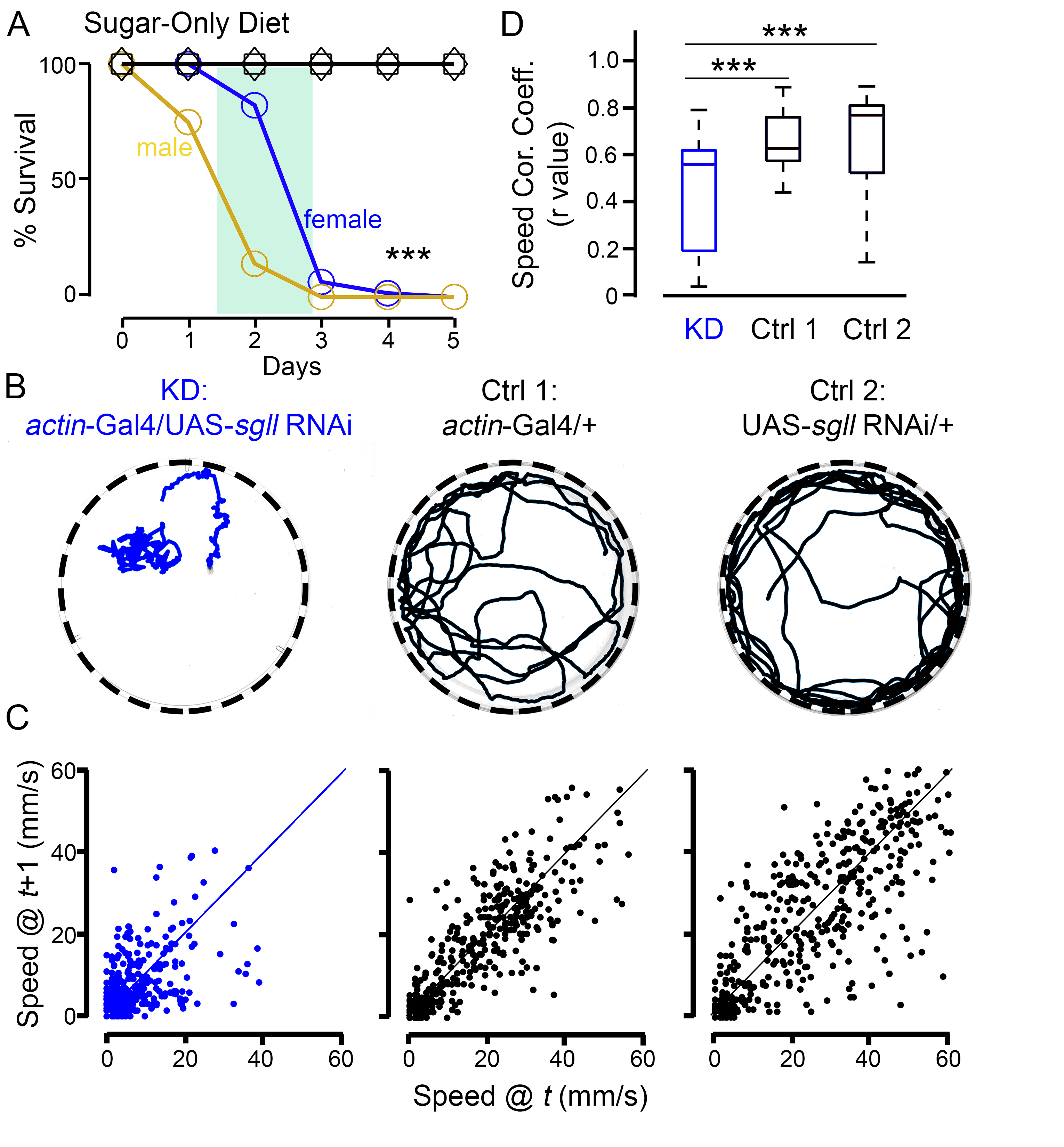

Figure S1. **Ubiquitous *sgll* KD flies show lethality and seizure-like behavior when reared on the sugar-only diet**. A) Survival of KD ­(blue and golden) and parental control flies (black square and black diamond) on the sugar-only diet. The green block indicates the time window in which the behavioral characterization was performed. *** *P* < 0.001, Log-rank test, n = 56-60 per genotype per sex. B) Representative travel traces from video recordings. C) Speed correlation plots between frame *t* and frame *t*+1. Each plot is corresponding to the one in panel B. D) Summary plot of speed correlation coefficient (Speed Cor. Coeff.). *** *P* < 0.001, One-way ANOVA with Tukey’s *post hoc*, n = 27-34 per genotype per condition.
