## Supplemental Figure 2 for "Pyridox(am)ine 5’-phosphate oxidase deficiency induces seizures in *Drosophila melanogaster*"

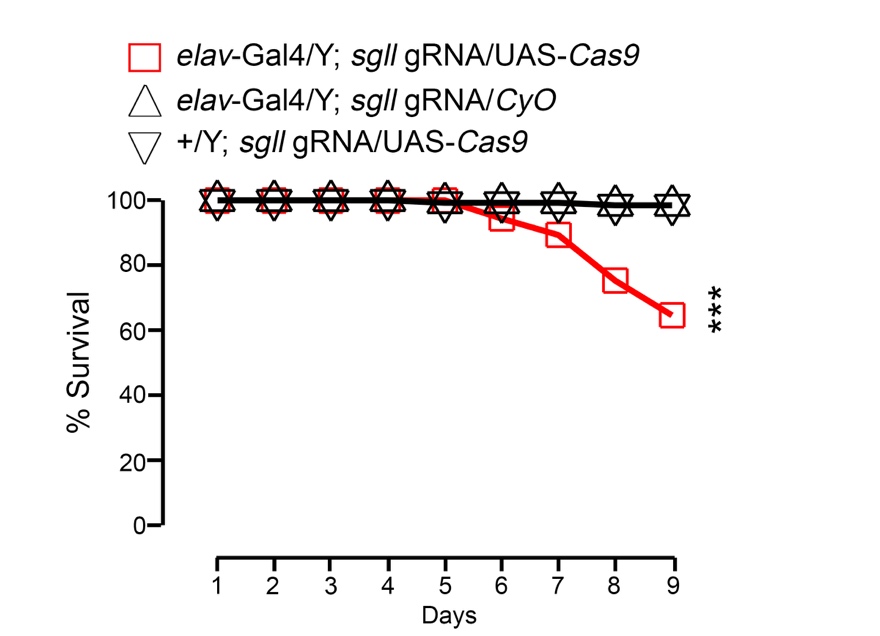

Figure S2. **Neural-specific *sgll* KO flies show lethality when reared on the sugar-only diet**. Survival of neural-specific *sgll* KO (genotype: *elav*-Gal4/Y; *sgll* gRNA/UAS-*Cas9*, red square) and control flies (genotypes: *elav*-Gal4/Y; *sgll* gRNA/*CyO* and +/Y; *sgll* gRNA/UAS-*Cas9*, black triangles) on the sugar-only diet. *** *P* < 0.001, Log-rank test, compared to control lines, n = 133-313 per genotype.
